# On the Number of Siblings and *p*-th Cousins in a Large Population Sample

**DOI:** 10.1101/145599

**Authors:** Vladimir Shchur, Rasmus Nielsen

**Author notes:** Corresponding author is V.Shchur.

## Abstract

The number of individuals in a random sample with close relatives in the sample is a quantity of interest when designing Genome Wide Association Studies (GWAS) and other cohort based genetic, and non-genetic, studies. In this paper, we develop expressions for the distribution and expectation of the number of *p*-th cousins in a sample from a population of size *N* under two diploid Wright-Fisher models. We also develop simple asymptotic expressions for large values of *N*. For example, the expected proportion of individuals with at least one *p*-th cousin in a sample of *K* individuals, for a diploid dioecious Wright-Fisher model, is approximately 1 − *e*^−(2^2*p*−1^)*K/N*^. Our results show that a substantial fraction of individuals in the sample will have at least a second cousin if the sampling fraction (*K/N*) is on the order of 10^−2^. This confirms that, for large cohort samples, relatedness among individuals cannot easily be ignored.

## 2. Introduction

As genomic sequencing and genotyping techniques are becoming cheaper, the data sets analysed in genomic studies are becoming larger. With an increase in the proportion of individuals in the population sampled, we might also expect an increase in the proportion of related individuals in the sample. For example, Moltke *et al*. (2014) found in a sample of 2,000 Inuit from Greenland that almost half of the sample had one or more close relatives in the sample. The census population size for Greenland Inuit is only about 60,000 individuals and the effective population size might be substantially lower. Henn *et al*. (2012) found 5000 pairs of third-cousin and 30,000 pairs of fourth cousin relatives in a sample of 5000 selfreported Europeans, with nearly every individual having a detected cryptic relationship. In Genome Wide Association Studies (GWAS), related individuals are routinely removed from the sample, but other strategies also exist for using relatedness as a covariate in the statistical analyses (e.g., Visscher *et al*. 2008). These observations raise the following question: given a particular effective population size, how many close relatives would we expect to find in a sample? The answer to this question may help guide study designs and strategies for addressing relatedness in population samples and improve design for GWAS. Of particular interest is the number of individuals in the sample without relatives, i.e. the number of individuals remaining in the sample if individuals with relatives are removed.

Substantial progress has been made on understanding the structure of a pedigree in a population. For example, Chang (1999) showed that the most recent common ancestor of all present-day individuals is expected to have lived *log*_2_(*N*) generations in the past if *N* is the population size. A great deal of progress has also been made in understanding the difference between genealogical processes in full diploid pedigree models versus the approximating coalescent process (e.g., Wakeley et al. 2012; Wilton et al. 2016). However, the distribution and expectation of the number of individuals with relatives in a random population sample is still unknown.

In this paper we will address this question by exploring two diploid and dioecious Wright-Fisher models. We will use these models to derive distributions and expectations of the number of individuals that have, or do not have, siblings, first, second, etc. cousins within a sample.

## 3. Dioecious Wright-Fisher Model

The Wright-Fisher model (Fisher 1930; Wright 1931) describes the genealogy of a population with constant effective population size *N*. The model assumes that generations do not overlap. Let 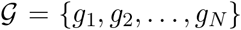 and 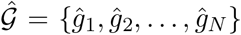 be two successive generations with *N* individuals in each. Then for each individual *ĝ_i_* from 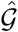 a parent *g_j_* is selected randomly and uniformly from 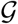.

In our study we consider a diploid population where each individual has two parents, one male and one female. Similarly to the original haploid Wright-Fisher model, the dioecious Wright-Fisher model (see e.g. Nagylaki 1997, King *et al*. 2017) assumes that generations do not overlap and, for each individual, the parents are chosen from the previous generation uniformly at random. The difference is that instead of a single parent, in the dioecious case, each individual has two parents, one male and one female, which are drawn independently from the corresponding sets of males and females in the preceding generation. We will refer to this model as the ‘non-monogamous Wright-Fisher model’ because we will also consider a model in which female and male parents form monogamous pairs. We will refer to the latter model as the ‘monogamous Wright-Fisher model’. As we will assume exactly equal proportions of males and females, the monogamous Wright-Fisher model is identical to the bi-parental monoecious model in King *et al*. (2017).

For both the non-monogamous and monogamous models, we assume that there are exactly *N* male and *N* female individuals. Each individual from generation 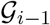 (we enumerate generations backward in time starting from 0, i.e. 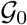 is the present generation and 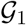 is the generation of parents of individuals from 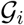) is assigned to a parent pair (one male and one female parent) from 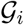. As we described above, under the non-monogamous model, male and female parents are chosen independently from each other for every individual. In the monogamous case, the parent pairs are fixed, i.e. we assume each male and female is part of exactly one potential parent pair.

The two diploid models are similar to each other in that the marginal distribution of the number of offspring of each individual is binomially distributed with mean 2. However, they differ from each other in the correlation structure among parents. The important difference between these two models is that the monogamous model does not allow for half-siblings (we say that two individuals are half-siblings if they share only one parent). On the other hand under the non-monogamous model, full siblings (individuals which share both parents) have a very low probability of appearing.

We note that other dioecious versions of the Wright-Fisher models could be considered with varying degree of promiscuity, but most would likely have distributions of relatedness that are somewhat intermediate between these two models, as long as they otherwise maintain Wright-Fisher dynamics. We also note that none of these models probably accurately describe the behaviour of human populations, which likely have a much higher variance in offspring number, variable population sizes, etc.

As mentioned above, individuals are siblings if they have the same parents. If individuals share only one parent, we call them half-siblings. We say that two individuals are *p*-th cousins if there is at least one coalescence between their genealogies in generation 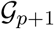. Of course, the amount of shared genetic material would depend on the number of shared ancestors in a certain generation. For two individuals, the number of shared ancestors is given in the supplementary materials of King *et al*. (2017) (see the discussion below). Notice, that two individuals can have different relations simultaneously. An example of such a situation is given in Figure 1: the individuals related by this genealogy are half-siblings and first-cousins at the same time.

**Figure 1.**
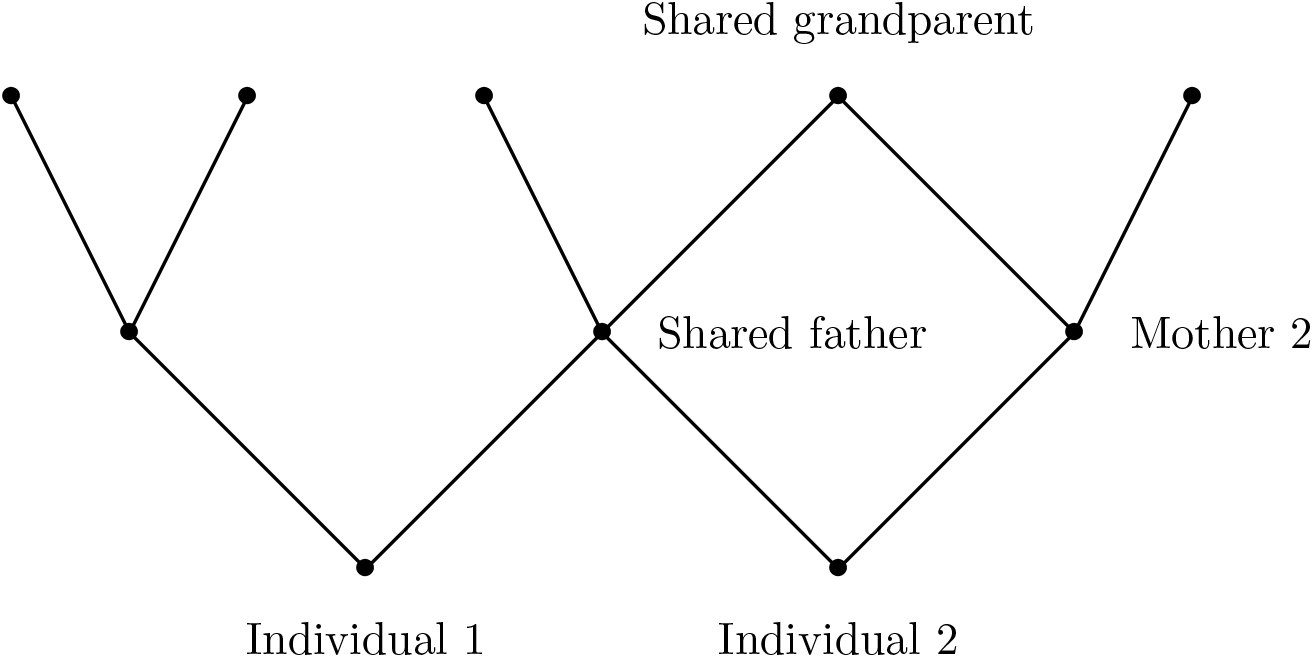
The two offspring (Individual 1 and Individual 2) related by this genealogy are half-siblings and first cousins at the same time. Notice that Individual 1 has a tree-like genealogy (no cycles, no inbreeding). The second individual though has inbreeding in its genealogy.

Let 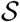 be a random sample of size *K* of individuals from the present-day generation 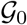 of a population described by either a monogamous or non-monogamous Wright-Fisher models. In this paper we derive the number *U_T_* (notation for monogamous case) or *V_T_* (notation for non-monogamous case) of individuals in 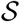 which do not have (*T* – 1)-order cousins (*T* = 1 would stand for (half-)siblings, *T* = 2 for first cousins, etc.) within 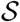 and have genealogy with no cycles. We will derive the probability distribution of *U*_1_ and *V*_1_ and expectations of *U_T_* and *V_T_* for *T* ≥ 2 in terms of Stirling numbers of the second kind. Further we present a simple analytical approximation of expectations of *U_T_* and *V_T_*. We derive this approximation as an exponential function of the ratio of the sample size to the effective population size.

The condition that individual’s genealogy does not have cycles means that there is no inbreeding in the history of the individual. Indeed, a cycle appears when two mating individuals share an ancestor, hence they are related to each other. On the contrary, if there is no inbreeding within *T* generations of ancestors of a certain individual, then all the ancestors have different parents, hence in the 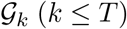 there are exactly 2^*k*^ ancestors of the individual under consideration.

Notice that the requirement that there is no inbreeding is satisfied as long as 2^*T*^ is small compared to the effective population size *N*. In this paper we are particularly interested in large populations. We will compute the fraction of individuals with siblings (*T* = 1) or *p*-th cousins (*T* = *p* + 1) in a sample in the limit of the effective population size *N* going to infinity. For fixed values of *T* and the sample size, *K*, the number of siblings and cousins goes to zero in the limit of large *N*. However, for a fixed ratio *K*/*N*, there is a positive expected number of siblings and offspring, but the expected number of cycles in the genealogy is small compared to *K*. This observation follows from the fact that the probability that two individuals share a parent is 1/*N*, which is a rare event for large *N*. Hence for large *N* all the ancestors of an individual are unrelated with high probability. We will, therefore, approximate the number of individuals who have siblings (or *p*-th cousins) by *K* – *U_T_* or *K* – *V_T_* depending on the model. We notice that using this method we cannot characterise, for example, the overlap between the set of individuals who have siblings and the set of individuals who have first-cousins, so we cannot provide an approximation of the number of individuals who have at least some kind of relatives within several generations.

Every genealogy has the same probability under the model. Hence our problem is equivalent to counting the number of possible genealogies with certain properties. To enumerate different genealogies, we will use the following approach. Firstly, we divide a sample 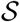 into subsets of siblings (in case of non-monogamous model, we create two independent partitions of the sample, one of partitions corresponding to shared fathers and the other corresponding to shared mothers). Then we assume that individuals from the same subset have the same parent couple (in the case of the monogamous model) or the same father or mother (in the case of the non-monogamous model), and individuals from different subsets have different parents. This approach is the basis for our analyses and leads us to the proof of formulas for expectations of *U_T_* and *V_T_*.

The combinatorial technique used to obtain exact formulas for expectations of *U_T_* and *V_T_* is very similar to the technique used in King *et al*. (2017) (see supplementary materials S1). In particular, we have to keep track of the number of ancestors at each generation which is the question of interest of the section S1.1 of King *et al*. (2017). Notice, that results in our paper and the result of S1.2 of King *et al*. (2017) complement each other. We find the expected number of individuals in a sample which do not have any relatives with respect to a certain generation, hence we know approximately the number of individuals which share at least one ancestor in that generation with at least one more individual from the given sample. However we cannot characterise finer relatedness (e.g. the number of shared ancestors in a given generation,) as more than one coalescence per generation between genealogies of two individuals is possible. The pairwise analysis of individuals can be performed using King *et al*. (2017) results, though it can be computationally challenging. The asymptotic behaviour derivation for *E*(*U_T_*)/*K* and *E*(*V_T_*)/*K* (for fixed *K*/*N* ratio) is a completely new result to the best of our knowledge.

We remind the reader that the Stirling number of the second kind *S*(*n, k*) is the number of ways to partition a set of size n into *k* non-empty disjoint subsets. A generalisation of this is the *r*–associated Stirling number of the second kind, *S_r_*(*n, k*) (Comtet 1974), which is the number of partitions of a set of size *n* into *k* non-empty subsets of size at least *r*. We provide more detailed information on the Stirling numbers of the second kind in the Appendix.

## 4. Probability distribution *U*_1_

We say that two individuals are siblings if they have the same parents. In this section we study the number of individuals *U*_1_ without siblings within a sample of a population. We derive both the probability distribution and expectation of *U*_1_.

### Theorem 1.

Let *U*_1_ be a random variable representing the number of individuals in a sample 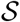 of size *K* without siblings in 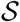 under monogamous dioecious Wright-Fisher model. Then

- the probability distribution of *U*_1_ is

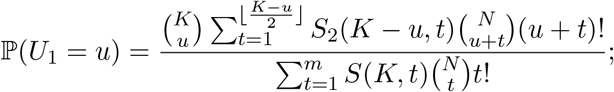
- the expectation of *U*_1_ is

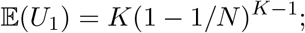
- if *K/N* = *α*

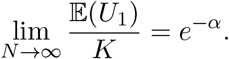

*Proof*. We begin the proof by computing the number of possible partitions of 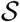 into *u* subsets of size 1 and *t* subsets of size greater than or equal to 2. Each subset of such a partition corresponds to the descendants in 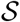 of the same couple of parents from 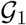. There are 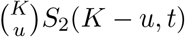 such partitions (see figure 2). Here the first multiplier corresponds to the number of choices of the first *u* individuals and the second multiplier corresponds to the number of partitions of the remaining *K* – *u* individuals into *t* disjoint subsets.

Now we need to assign *u* + *t* subsets to different couples of parents from 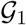. There are 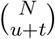 possibilities for choosing couples that have descendants in 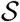 and (*u*+*t*)! permutations which assign these particular couples to different subsets of the given partitions of 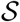.

Finally, summing over all possible values of *t* we get

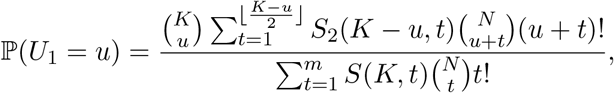

where ⌊·⌋ stands for the floor integer part.

**Figure 2.**
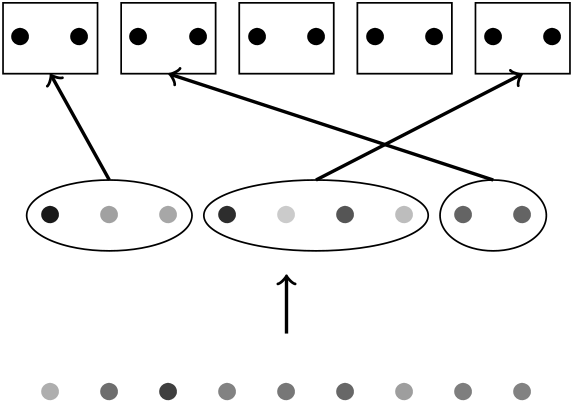
Illustration to the proof of Theorem 1. Each dot correspond to an individual. The bottom set of points corresponds to the individuals in the sample 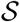. This sample is divided in disjoint subsets (the set of points in the middle): this partition corresponds to sets of siblings, or in other words individuals from each subset will be assigned to the same couple of parents. The top row corresponds to the set of couples in the parent generation. Subsets of siblings (from the middle row) are assigned to different couples of parents (from the top row).

The expression for expectation of *U*_1_ is much simpler. The probability *π*_1_ that an individual 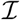 does not have any siblings in 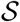 is *π*_1_ = (1 – 1/*N*)^*K*−1^, because all other individuals from 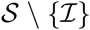 can be assigned to any couple of parents except for the parents of the individual 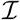. By linearity, the expectation of *U*_1_ is

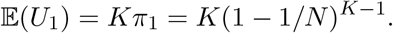

To prove the last statement of the theorem it is enough to rewrite

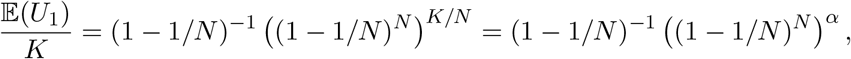

because *K/N* = *α* by definition. Now notice that

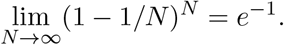

Hence the last statement of the theorem is proved

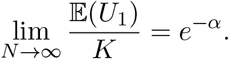

## 5. Expectation of *U*_2_

In this section we will provide an expression for expectation of the number *U*_2_ of individuals in a sample which do not have first cousins in this sample. We will also establish a limit for 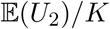 in the case of a fixed ratio between *K* and *N*.

### Theorem 2.

Let *U*_2_ be a random variable representing the number of individuals in a sample 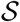 *of size K without first cousins in* 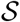 under a monogamous dioecious Wright-Fisher model. Then the expectation of *U*_2_ is

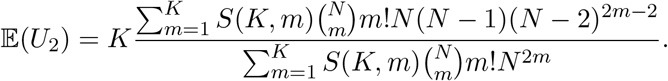

*Proof*. Similarly to the case of 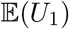, we need to find the probability *π*_2_ for a single individual not to have first cousins within 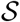. Then the expectation 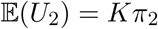. Denote individuals from 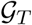 which have descendants in 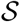 by 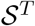.

Choose an individual 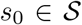, let 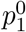 and 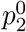, be parents of *s*_0_. If *s*_0_ does not have first cousins, then 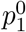 and 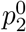 are assigned to different couples from 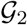 and those couples do not have other descendants in 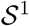.

Similarly to derivation of distribution of *U*_1_, we first partition 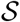 into *m* disjoint subsets. We choose *m* couples from 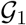 and establish a one-to-one correspondence between the subsets and the couples. There are *N* possibilities to choose a couple of parents for 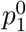, *N* – 1 choices for 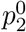 and (*N* – 2) choices for all other 2*m* – 2 individuals from 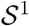. Summing over *m* we get

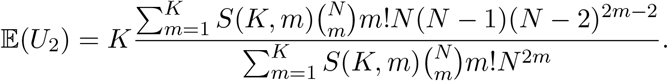

Our next goal is to find the limit of 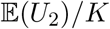 for a fixed ratio of sample size to the population size. We assume that *K/N* = *α* for some constant 0 ≤ *α* ≤ 1 and we consider the limit of 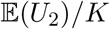 for *K* → ∞.

### Theorem 3.

Let 0 ≤ *α* ≤ 1 and set *K* = *αN*. Then

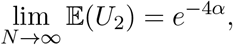

The following lemma states that the sum of the first *βK* terms of the series in the formula for 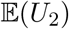 is small for large values of *K*. This makes it possible to make further approximations under the hypothesis that *m* = *O*(*K*).

### Lemma 1.

Let *K* = *αN* for some 0 ≤ *α* ≤ 1 and set *β* = (2 ln 2)^−1^. Then

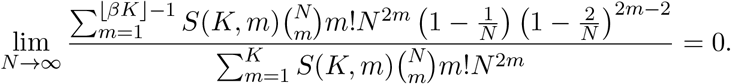

*Proof*. Denote

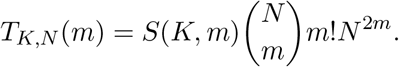

First, notice that

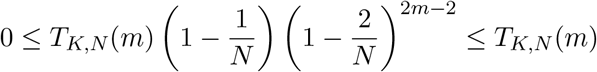

We will show that for *β* = (2 ln2)^−1^ < 1/2

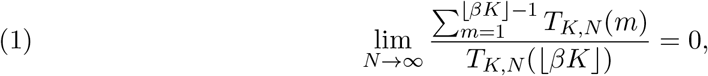

which will immediately prove the statement of the Lemma.

Our goal is to prove that

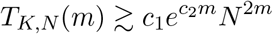

for some constants *c*_1_,*c*_2_ and *K* large enough.

We begin by approximating the following ratio for *m* ≤ ⌊*βK*⌋

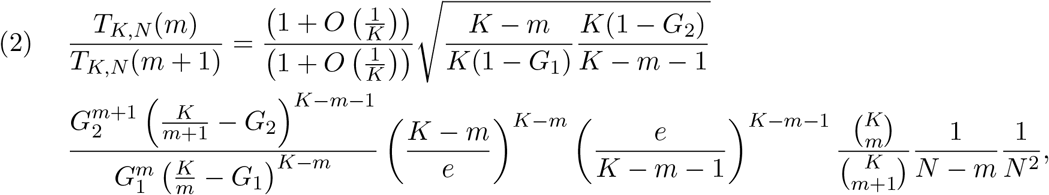

by applying approximation (11). Here *G*_1_ = *G*(*K, m*) and *G*_2_ = *G*(*K, m* + 1).

Notice that 0 < *G*_1_ < *G*_2_ < − *W*_0_(−2*e*^−2^) < 1/2. The following term is bounded by a constant (we remind the reader that 0 < *m* ≤ *βK* < *K*/2)

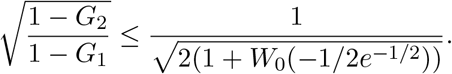

After simplification, all the factorials in the formula are of the form (*constK*)!, hence they can be approximated uniformly in *K* by Stirling’s approximation

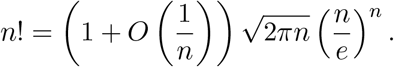

For simplicity of notations we drop all terms 1 + *O*(1/*K*) in (2). We also notice that

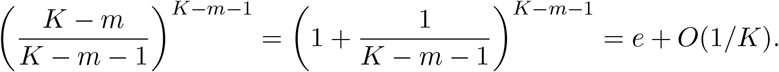

So for *K* large enough the ratio (2) has the following approximation

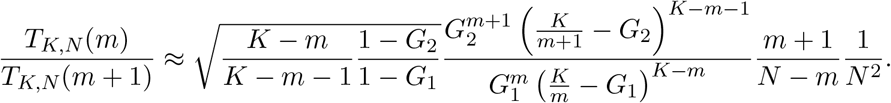

The derivative of *G*(*x*)^1/*x*^(*x* – *G*(*x*))^1−1/*x*^ (*x* ≥ 1) with respect to *x* is

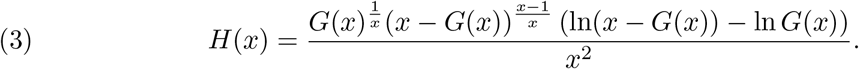

*H*(*x*) has one real root *x* = 2ln2 if *x* ≥ 1. The derivative *H*(*x*) is positive for *x* > 2ln2, so *G*(*x*)^1/*x*^(*x* – *G*(*x*))^1−1/*x*^ is an increasing function of *x* for *x* > 2 ln 2. Hence as soon as *K/m* > 2ln2, or *m* < *K*/(2ln2), the following inequality holds

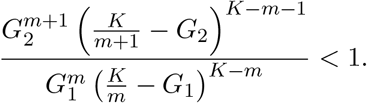

Consequently, for sufficiently large *K* we obtain the following upper bound for (2)

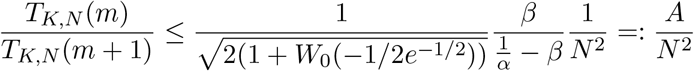

Hence, by recursion for *m* < ⌊*βK*⌋

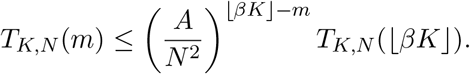

Now we use the obtained inequality to prove limit (1)

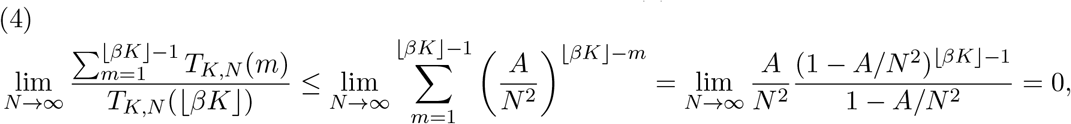

where the second equality holds by summing over the geometric progression.

### Lemma 2.

Let *K* = *αN* for some 0 ≤ *α* ≤ 1, set *β* = (2 ln 2)^−1^. Then for any *m* such that ⌊*βK*⌋ ≤ *m* < *K*

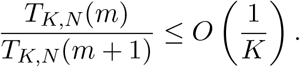

*Proof*. From the proof of Lemma 1, for *K* large enough and for *β* ≤ *m/K* ≤ 1

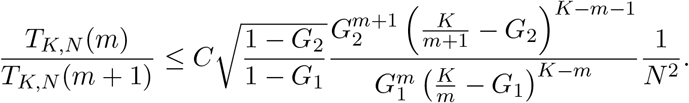

Notice that *xe^x^* = −1/*e* + *O*((*x* − 1)^2^) near *x* = −1. Hence 1 − *G*(*x*) = *O*(|*x* − 1|) and *x* − *G*(*x*) = *O*(|*x* − 1|) for *x* → 1. By definition, the Lambert *W*-function (Olver *et al*. (2010)) is the inverse function of *xe^x^*. If *x*_1_ > −1 and *x*_2_ < −1 are two points in the neighbourhood of −1 such that *x*_1_*e*^*x*1^ = *x*_2_*e*^*x*2^, then |*x*_1_ – *x*_2_| = *O*(|*x*_1_ – 1|) = *O*(|*x*_2_ – 1|). For *x* > 1, −*xe*^−*x*^ ∈ [−1/*e*; 0]. The value of the main branch, *W*_0_(*xe^x^*), is in the interval [−1,0]. So −*x* and *W*_0_(−*xe*^−*x*^) correspond to *x*_1_ and *x*_2_.

Hence

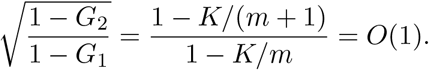

Now we use mean value theorem to approximate

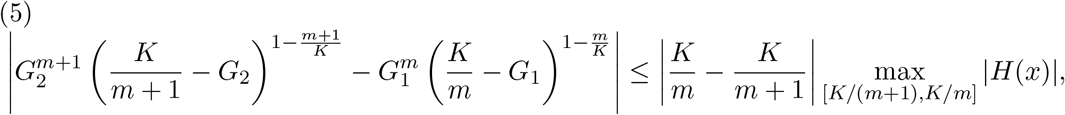

where *H*(*x*) is given by expression (3). Denote Δ*x* = |*x* – 1|, and notice that

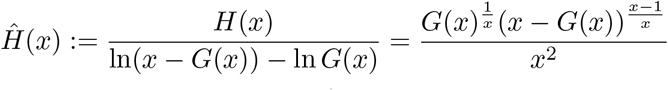

and ln *G*(*x*) are continuous near *x* = 1 and *Ĥ*(1) = 1, ln *G*(1) = 0. So for small Δ*x*

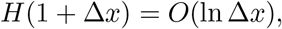

and hence

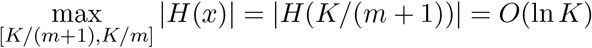

which leads to the approximation of (5) with *m* = *O*(*K*)

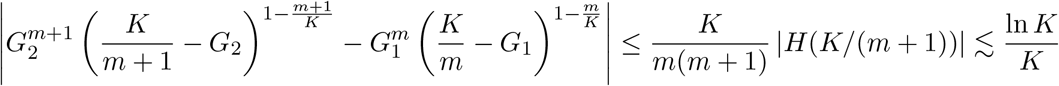

We use this estimate and the Taylor expansion of logarithm to get

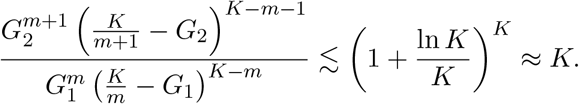

Finally, we estimate the ratio *T_K,N_*(*m*)/*T_K,N_*(*m* + 1) for *K* large enough

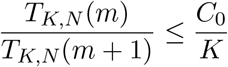

with some constant *C*_0_, which depend on *α*.

Now we are ready to prove the theorem.

*Proof*. Firstly, notice that

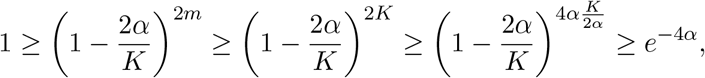

and (1 – 1/*N*)(1 – 2/*N*)^2^ → 1 as *N* → ∞. Hence, the lower bound is valid for any *α* and *K*

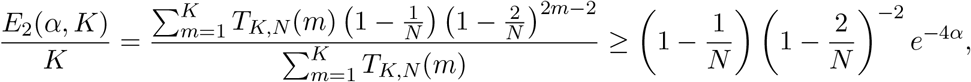

where the right part trivially converges to *e*^−4*α*^ with *N* → ∞ (we remind that *K* = *αN* for some constant 0 ≤ *α* ≤ 1).

Now we prove that this bound is sharp by applying subsequently Lemmas 1 and 2

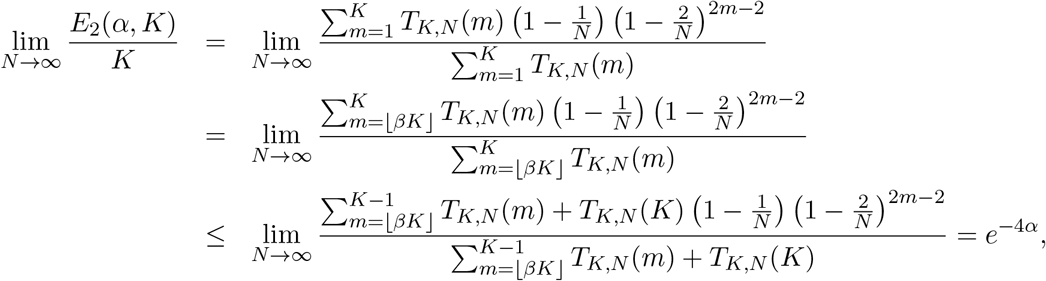

because from Lemma 2 it follows

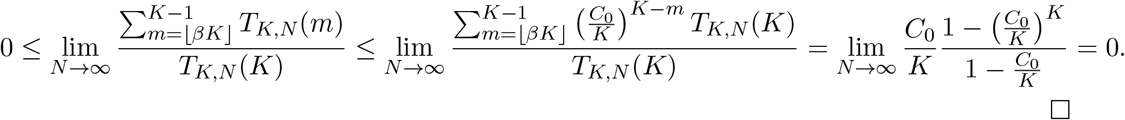

## 6. General Case: Expectation of *U_p_* for *p* ≥ 2

Similarly to the expectation of *U*_2_, we can find the probability of the expected numbers *U_p_*(*p* ≥ 2) of individuals which do not have (*p* − 1)-cousins and with pedigrees without cycles.

### Lemma 3.

Let 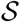 be a set and 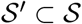 be a subset of size 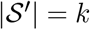. The number of partitions of a set 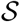 of size *N* into *M* disjoint subsets such that all elements of 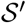 are in different subsets is

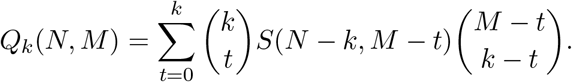

*Proof*. Let 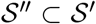, 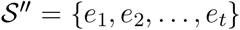, such that each element, 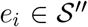, makes its own subset *P_i_* = {*e_i_*} in the partition of 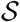. If 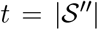 there are 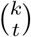 ways to choose such a subset. Then, 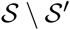 should be split into *M* – *t* non-empty subsets, *P*_*t*+1_, *P*_*t*+2_,…, *P_M_*, to obtain a partition of 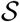 into exactly *M* subsets. There are *S*(*N* – *k, M* – *t*) possible ways of doing that. Each of the *k* – *t* elements of 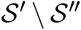 are then added to distinct subsets among the remaining *M* – *t* subsets, *P_i_, i* > *t*, which can be done in 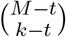 ways.

Summing over all possible values of t we prove the statement.

*Remark 1*. For *k* = 1, Lemma 3 turns into the well-known recursive formula for Stirling numbers of the second kind.

The next theorem establishes the expression for the expectation of *U_p_* and its limit for fixed *K* to *N* ratio in the general case. Due to the size of the formula we had to introduce additional notations for readability.

### Theorem 4.

- For any natural *p ≥ 1* the expectation of *U_p_* is

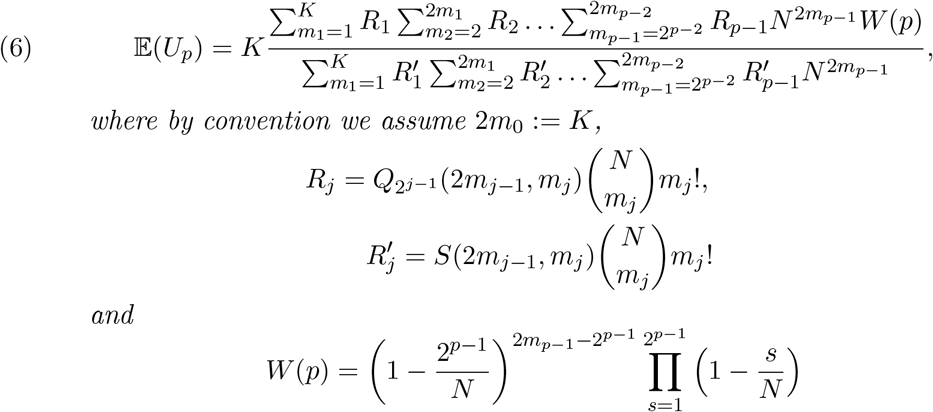

and

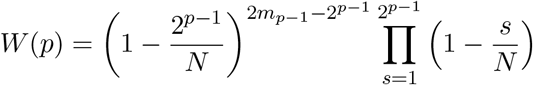
- If *K* = *αN* (*i* = 1, 2,…,*p*), then

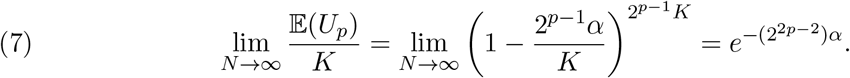

*Proof*. To prove the first statement, we apply repeatedly the same arguments as used for Theorem 2: for each generation, we split the ancestors of the sample into subsets of siblings while controlling that ancestors of the given individual are not in the same subsets.

The proof of (7) is similar to the proof of Theorem 3. First we can show that we can substitute summations over *m_i_* > *βK* for some constant *β* (see Lemma 1). Then we use estimations for *Q_i_* that are similar to those obtained in Lemma 2.

## 7. Non-monogamous Wright-Fisher model

Similar results to those obtained for the monogamous case also hold for the non-monogamous dioecious Wright-Fisher model. However, in contrast to the monogamous case, the probability that two individuals are full siblings or full *p*-th cousins (i.e. sharing two ancestors) is rather small. Most familial relationships would involve sharing only one common ancestor at a given generation, i.e. related individuals would typically be half siblings or half *p*-th cousins.

Let *V_p_* be a random variable representing the number of individuals in a sample 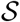 of size *K* without half siblings or full siblings (*p* = 1) or half *p*-th cousins or full *p*-th cousins (*p* ≥ 2) in 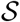 under the non-monogamous Wright-Fisher model. The next theorem established the expression for the expectation of *V_p_* and its limit for *K* → ∞ in the case of fixed ratio between *K* and the population sizes *N*.

### Theorem 5.

- For any natural *p* ≥ 1, the expectation of *V_p_* is

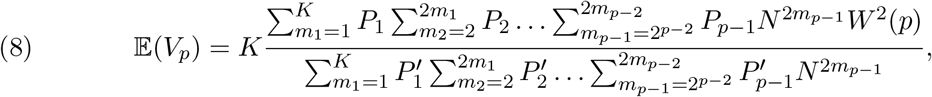

where we assume *m*_0_ = *K* and

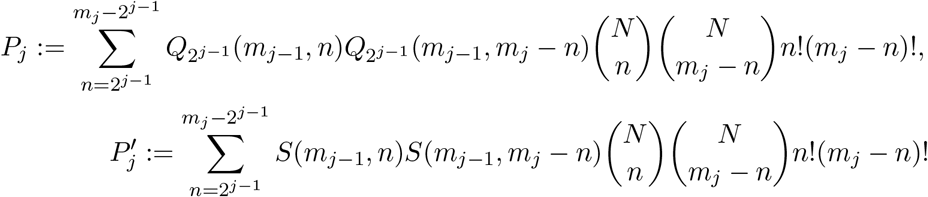

and

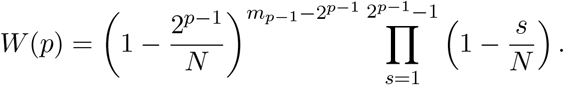
- If population sizes *K* = *αN*, then

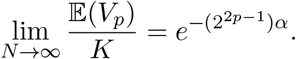 The proof of the theorem is similar to the case of the monogamous model. The function *P_j_* counts the number of possibilities to have exactly *m_j_* parents (male plus female) In particular,

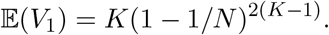

### Corollary 1.

The qualitative behaviour of *U_i_* and *V_i_* is the same, more precisely

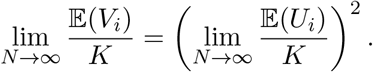

## 8. Numerical Results

In this section we present numerical results for expectations of *U_p_* and *V_p_, p* = 1,2, 3. Every plot of figures 3 and 4 represents the behaviour of 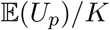 or 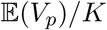 for a particular *p* = 1, 2, 3. Those values are computed by formulas (6) or (8) for different values of *N*(*N* = 20,100,200) as a function of the ratio *K/N*. We also add corresponding limiting distribution to every plot to illustrate the convergence.

**Figure 3.**
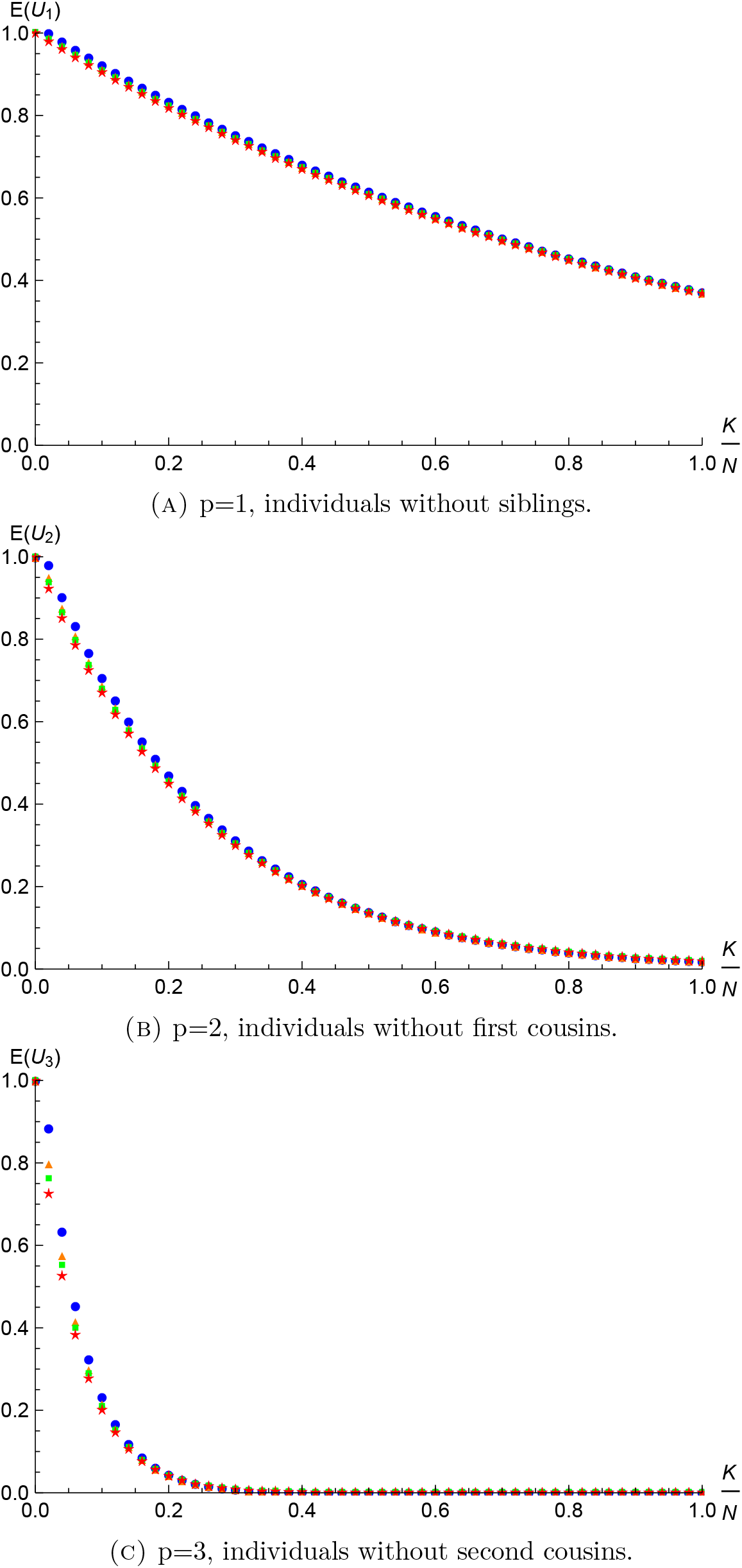
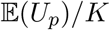 as a function of the *K/N* ratio for *N* = 50 (●), 100 (▲), 200 (▀) and the corresponding limiting distribution (⋆).

**Figure 4.**
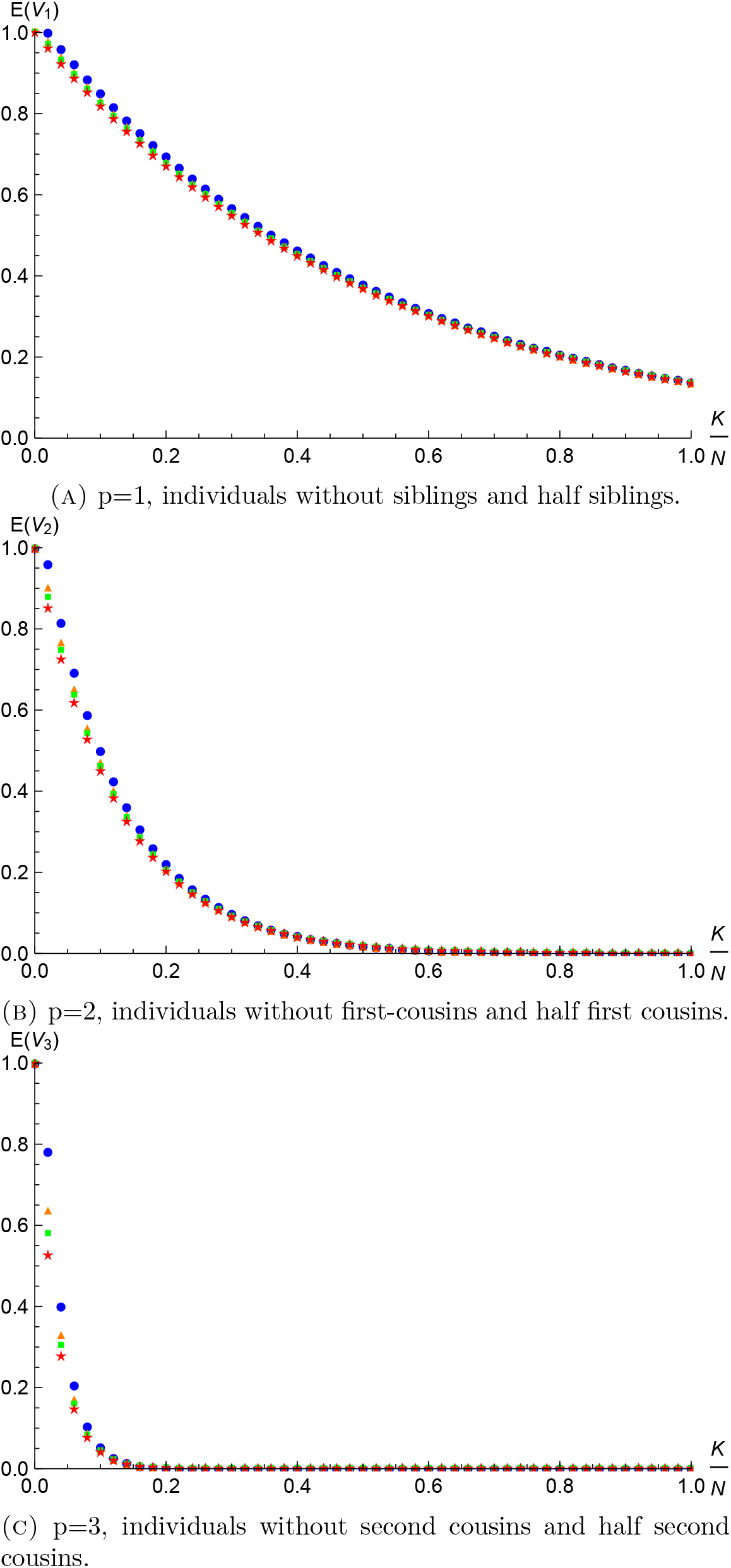
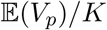 as a function of the *K/N* ratio for *N* = 50 (●), 100 (▲), 200 (▀) and the corresponding limiting distribution (⋆).

Because the effective population sizes are typically rather large (at least thousands of individuals) we might expect a satisfactory approximation of *E*(*U_p_*) and *E*(*V_p_*) by its limiting distribution even for relatively small *K/N* ratios. One can also check that in our proofs the errors in the estimates are of the order of 1/*N*, hence for the desired ratio we can estimate the absolute error for smaller values of *K, N* numerically and then increase *N* to get the desired precision.

## 9. Discussion

In this paper we analysed the expected values of the number of individuals without siblings and *p*-th cousins in a large sample of a population. To do that we used two extensions of Wright-Fisher model which keeps track of the two parents of an individual.

The first extension corresponds to a monogamous population and the second to a non-monogamous population. The two models represent two extremes in terms of degree of promiscuity, and we might expect that in most other dioecious versions of the Wright-Fisher model, with intermediate degrees of promiscuity, the number of individuals without siblings or *p*-th cousins is somewhere in between those two regimes - as long as the models otherwise maintain Wright-Fisher dynamics.

Under both models we derived expressions for these expectations under the hypothesis that the pedigrees have no cycles (except for the one appearing in full sibs). Notice that this restriction is not too strong, because one can easily show that the chance that an individual has a pedigree with a cycle is a second-order effect as soon as the number of ancestors (≤ 2^*p*^) in a generation is much smaller than the effective population size *N*.

The important result of the paper is the limiting distributions for 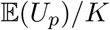 and 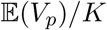. It turns out that 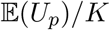 and 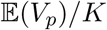 converge point-wise to *e*^−*cK/N*^ where the constant *c* is 2^2*p*−2^ for *U_p_* and 2^2*p*−1^ for *V_p_*.

We notice that even when the sampling fraction is relative low, the proportion of individuals in the sample with no close relatives can be small. For example, for the non-monogamous model and a sampling faction of 5%, the proportion of individuals with at least a second cousin is approx. 70% if the population size is at least *N* = 200. For a sampling fraction of 2% the proportion in individuals with at least a second cousin is close to 50% for reasonably large population sizes in case of random mating population or almost 30% in case of monogamous population. For sampling fractions on the order of 0.01 or larger, we expect a large proportion of individuals to have at least one other individual in the sample to which they are closely related. This fact should be taken into account in all genetic, and non-genetic, epidemiological studies working on large cohorts.

In the study of Danish population structure, Athanasiadis *et al*. (2016) discovered 3 pairs of first cousins and one pair of second cousins in a sample of just 406 individuals. Based on their estimate of an effective population size of 500,000, we would expect to find 1.32 individuals with first cousins under the monogamous model and 2.63 individuals with first cousins under the non-monogamous model. The empirical number of 3 first-cousins in the sample is therefore not significantly different of the expected number of 1.32 under the monogamous model assumption. It is also not statistically significantly different from the expected number of 2.63 under the non-monogamous model. The expected number of second cousins in the sample is 5.24 and 10.41 under the monogamous and non-monogamous models, respectively. The inferred number of 1 is much smaller than this, likely because it is difficult to infer second cousins empirically. We would in general expect that the true number of second cousins is larger than the true number of first cousins.

Notice, that the probability for two individuals to be *p*-th cousins is approximately 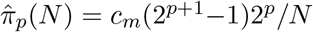, where *c_m_* is 1 for monogamous model and 2 for non-monogamous model. Hence, the expected number of pairs of *p*-th cousins in a sample of size *K* is approximately 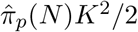. Henn *et al*. (2012) found approximately 5000 pairs of third cousins and 30000 pairs of fourth cousins in a sample of only 5000 individuals with European ancestry, which would be expected for effective population sizes of 2 · 10^5^ – 3 · 10^5^ under the monogamous model and twice that (4 · 10^5^ – 6 · 10^5^) under the non-monogamous model. These numbers are roughly compatible with estimates of effective population sizes obtained for modern European populations (e.g., Athanasiadis *et al*. (2016)). We note that effective population size is a tricky concept for a spatially distributed population such as European humans, but the breeding structure observed in these samples suggest that the degree of relatedness in the sample is compatible with population sizes on the order of 10^5^ – 10^6^.

## 10. Appendix: Stirling numbers of the second kind and their generalisation

In this section we provide definitions and properties of Stirling numbers of the second kind.

The Stirling number of a second kind *S*(*n,k*) is the number of ways to partition a set of size *n* into *k* non-empty disjoint subsets. These numbers can be computed using the recursion (Abramowitz and Stegun 1972)

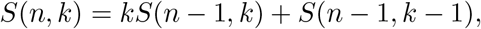

with *S*(0, 0) = *S*(*n*, 0) = *S*(0, *n*) = 0 for *n* > 0. Notice that *S*(*n, n*) = 1.

An *r*–associated Stirling number of the second kind, *S_r_*(*n, k*) (Comtet 1974), is the number of partitions of a set of size *n* into *k* non-empty subsets of size at least *r*. These numbers obey a recursion formula (Comtet 1974) similar to that for Stirling numbers of second kind

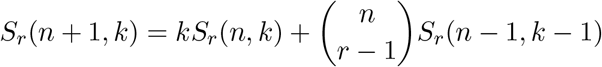

with *S_r_*(*n*, 0) = *S_r_*(1, 1) = 0. In particular, for *r* = 2

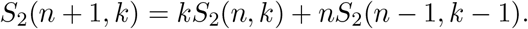

### 10.1 Uniformly valid approximation for *S*(*n, k*)

The following useful approximation of Stirling numbers of the second kind is established by Temme (1993)

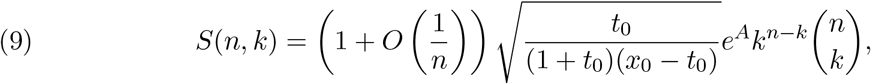

where *t*_0_ = *n/k* – 1, *x*_0_ ≠ 0 is the non-zero root of the equation

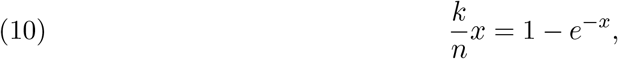

and

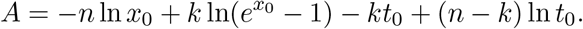

The following form of this approximation is known

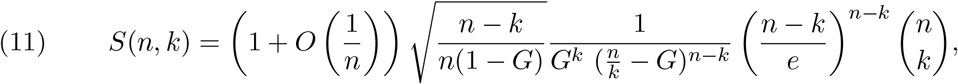

with *G* = −*W*_0_(−*n*/*ke*^−*n/k*^), where *W*_0_ is the main branch of Lambert *W*-function (Olver et al. 2010).

We did not find a reference for the formula (11) in the literature, so we provide briefly the proof. Notice that −1/*e* < −*n/ke*^−*n/k*^ < 0, hence *G* ∈ (0,1). Let us show that *x*_0_ = *n/k* – *G* is the non-zero root of equation (10)

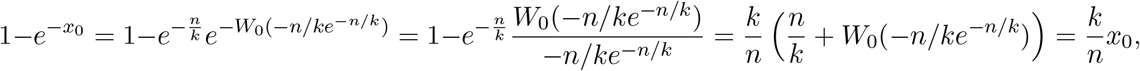

where the second equality is due to the Lambert function property *e*^−*W*(*x*)^ = *W*(*x*)/*x*. Substituting *t*_0_ and *x*_0_ in approximation (9) by their values and simplifying the formula, one gets the needed result. Obviously,

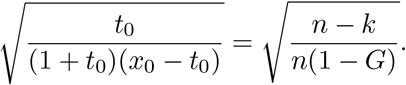

Now consider *e^A^k^n−k^*

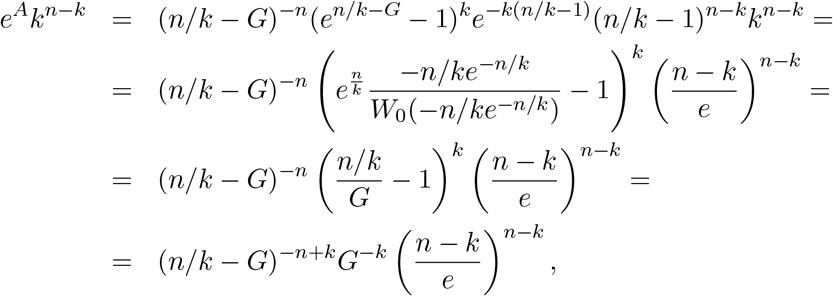

which finished the proof of equivalence of approximations (9) and (11).

## Acknowledgement

The work was supported by the UCOP Catalyst Award CA-16-376437.

